# Autonomous epithelial folding induced by an intracellular mechano–polarity feedback loop

**DOI:** 10.1101/2021.02.10.430574

**Authors:** Fu-Lai Wen, Chun Wai Kwan, Yu-Chiun Wang, Tatsuo Shibata

## Abstract

Epithelial tissues form folded structures during embryonic development and organogenesis. Whereas substantial efforts have been devoted to identifying mechanical and biochemical mechanisms that induce folding, how they interact remains poorly understood. Here we propose a mechano–biochemical model for dorsal fold formation in the early *Drosophila* embryo, an epithelial folding event induced by shifts of cell polarity. Based on experimentally observed apical domain homeostasis, we couple cell mechanics to polarity and find that mechanical changes following the initial polarity shifts alter cell geometry, which in turn influences the reaction-diffusion of polarity proteins, thus forming a feedback loop between mechanics and polarity. This model can induce spontaneous fold formation *in silico*, recapitulate polarity and shape changes observed *in vivo*, and confer robustness to tissue shape change against small fluctuations in mechanics and polarity. These findings reveal emergent properties of a developing epithelium under control of intracellular mechano–polarity coupling.

## I. INTRODUCTION

Epithelial tissues cover body surfaces and line internal organs and cavities in many metazoan animals. During development, epithelial cells undergo shape changes that orchestrate tissue-scale deformation such as convergent extension, branching and folding [11]. Epithelial cell shape changes typically require mechanical forces that are generated via biochemical processes that occur within the cells themselves, while these biochemical processes are governed by reaction, diffusion and transport of their molecular constituents, which may depend on the boundary conditions, including the cell shape [20]. Cell shape changes can thus influence the biochemical processes that induce these shape changes in the first place, thereby integrating mechanical forces and their underlying biochemical processes into a feedback loop [14]. Such a mechano–biochemical feedback loop has been proposed to play a pivotal role in coordinating cell morphogenesis and tissue reorganization during normal development as well as contributing to pathological conditions that involve dysregulation of tissue growth and cell movement [2, 10, 32].

Epithelial cells exhibit intrinsic polarity that allows them to perform specialized functions. Epithelial polarity can be established, and needs to be maintained, by a reaction-diffusion type mechanism that governs the non-uniform distribution of polarity regulators across the cell surface [5]. The partitioning-defective (PAR) polarity proteins display polarized distribution along the apical–basal axis of the cells to regulate the activity of spatially localized structural and signaling proteins, including those that produce, modulate and transmit mechanical forces that control epithelial deformation [24, 37]. Thus, modulation of the polarity system may affect cell mechanical properties, inducing cell shape changes that alter tissue architecture.

One such polarity-dependent deformation process is the formation of the epithelial folds that are generated on the dorsal side of the early fruit fly (*Drosophila melanogaster*) embryo, called dorsal fold formation [38, 40]. Distinct from the canonical epithelial folding events that require the motor protein myosin that contracts the actin cytoskeleton, *Drosophila* dorsal fold formation is initiated by the remodeling of the PAR-dependent epithelial polarity without changes of myosin activity [40]. Although experimental evidence suggests that differential modulation of PAR-dependent polarity between cells is essential for dorsal fold formation and that microtubule-based forces underlie polarity-dependent cell shape changes [38, 40], the exact mechanism by which the polarity remodeling process instigates the mechanical alteration to induce folding has not been resolved. Here we postulated that during dorsal fold formation cell shape changes impact the remodeling of epithelial polarity, thereby integrating cell deformation and polarity evolution into a feedback loop. We tested this hypothesis using mathematical modeling and verified our theoretical predictions via quantitative *in vivo* imaging of the *Drosophila* embryo. We show that such mechano–polarity feedback allows epithelial cells to autonomously change their shapes, leading to self-organized formation of epithelial folds. Our modeling results thus support the previously proposed hypothesis that external forces from the surroundings are not required for dorsal fold formation [40]. This model allows us to uncover self-organizing principles of tissue reorganization emerging from intracellular mechano–polarity coupling.

## II. MATHEMATICAL MODELING

To simulate the polarity-dependent dorsal fold formation in *Drosophila*, we constructed a three-dimensional (3D) model which consists of biochemical and mechanical modules as described below.

### A. The biochemical module for the formation of epithelial apical–basal polarity

The plasma membrane of the epithelial cells in the *Drosophila* blastoderm embryo are polarized along the apical–basal axis (Fig. 1A). The key polarity regulators, atypical protein kinase C (aPKC), Par-1, and Bazooka, are each localized to, and thereby respectively define, the apical and basolateral domains as well as the interface domain between these two [30]. To model epithelial polarity formation, we constructed a minimal reaction-diffusion model following a previous non-epithelial model developed for the one-cell stage *C. elegans* embryo [12]. Our three-element network model (”*A*–*B*–*P* network”; Fig. 1B) contains the apical and basal-lateral proteins, aPKC and Par-1 (denoted by *A* and *P*, respectively), as in the *C. elegans* model. In addition, we added a third and epithelium-specific element, Bazooka (denoted by *B*), that specifies the subapical region where cell-cell adherens junctions are positioned [16]. Since the reactions responsible for the polarity formation take place when the polarity regulators associate with the plasma membrane, we assumed that the polarity system can be approximated by a surface reaction that is coupled to a cytoplasmic pool of polarity regulators. Given that the membrane distribution of polarity regulators is approximately rotationally symmetric with respect to the apical–basal axis, we simplified the surface reaction by considering the spatial distribution of regulator concentrations along the apical–basal axis in one dimension.

**FIG. 1.**
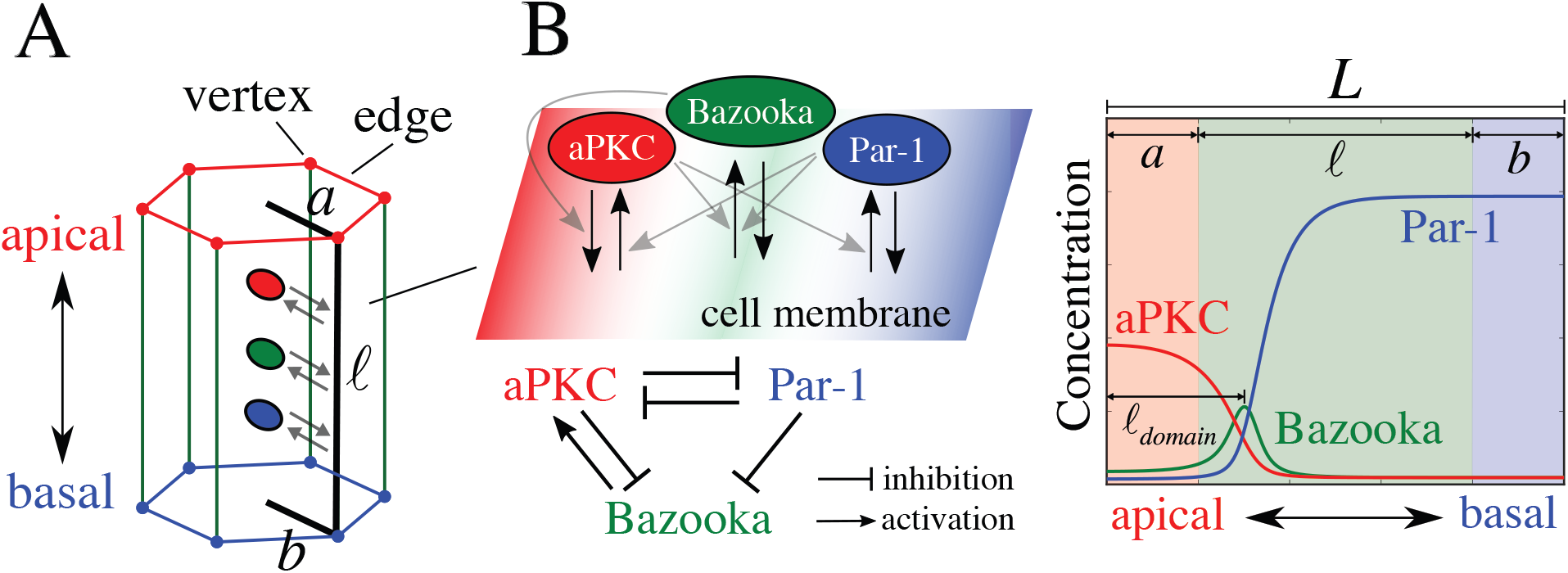
Schematics of the modeled epithelial cell and its polarity network. (A) A hexagonal prism depicting the geometry of the modeled epithelial cell, with its vertices and edges color coded to illustrate their distinct mechanical and biochemical properties. The arrows indicate the exchanges between the cytoplasmic and the membrane pools of the polarity proteins (shown as the color-coded ovals). (B) (Upper left) Enlarged view of the cytoplasm-membrane exchanges depicted in (A). Black arrows denote the association/dissociation of the polarity proteins to/from the membrane surface. The parallelogram with three distinct color regions for the three domains of the polarity regulators – aPKC, Bazooka, and Par-1 – depicts the membrane surface. Grey arrows denote the regulatory interactions among the three polarity regulators that modulate these association/dissociation exchanges. (Lower left) A diagram for the modeled polarity network with arrows depicting the activation/inhibition interactions. See Mathematical Modeling section for references that reported these interactions. (Right) Schematic representation of a 1D concentration profile along the apical–basal axis for the three polarity network components. The color-coded shades represent the apical, lateral, and basal regions of the cell membrane corresponding to the color-coded edges depicted in (A). See Mathematical Modeling section for the definition of *a*, *b* and *ℓ*.

The evolution of concentrations are thus formulated as follows,

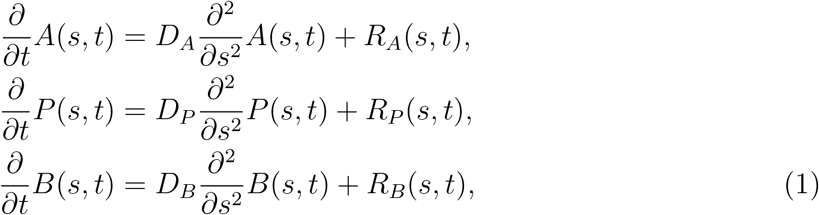

where *A*(*s, t*), *P* (*s, t*), and *B*(*s, t*) denote the membrane concentration of aPKC, Par-1, and Bazooka, respectively, at time *t* and membrane location *s* along the apical–basal axis (0 ≤ *s* ≤ *L*, see Fig. 1B). *D_A_*, *D_P_*, and *D_B_* are respective diffusion coefficients that govern the diffusion of these polarity regulators on the membrane surface.

The second terms in Eq. (1) describe the membrane association or dissociation reaction kinetics of the polarity regulators. These kinetics depend on the molecular interactions among the regulators themselves. Based on previously established polarity protein interactions in various epithelial systems [36], we consider phosphorylation-mediated mutual inhibition of the membrane association between aPKC and Par-1 [8, 21], the inhibition of membrane association of Bazooka by both aPKC [7, 39] and Par-1 [3], and the membrane recruitment of aPKC by Bazooka [25] (Fig. 1B, left). The reaction terms thus read,

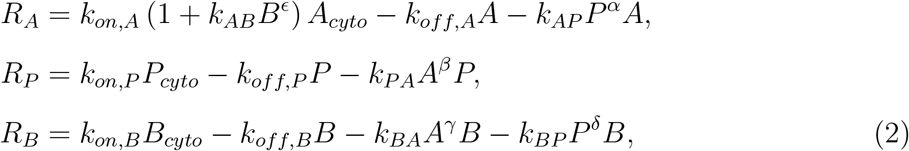

wherein *k_on/off,A_*, *k_on/off,P_* and *k_on/off,B_* are the membrane association (or dissociation) rates per unit concentrations in the cytosol (or on the membrane), *k_XY_* with *X, Y* ∈ {*A, P, B*} is the membrane dissociation rates resulted from the mutual antagonism between regulators *X* and *Y*, *k_AB_* is the membrane association rate of *A* due to the presence of *B*, whereas *α*, *β*, *γ*, *δ*, and *∊* are the parameters of these molecular interactions.

We assumed that the regulators diffuse much faster in the cytosol than on the cell membrane. The cytosolic concentration is thus taken to be uniform in space and can be calculated from the principle of mass conservation, as follows:

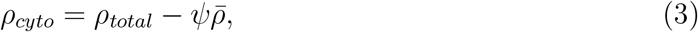

where *ρ_total_* is the total concentration (total number of molecules per unit volume of the cell) with *ρ* ∈ {*A, P, B*}, *ψ* is the surface-to-volume conversion factor (see below for details), while 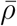 is the average concentration across the membrane, calculated as

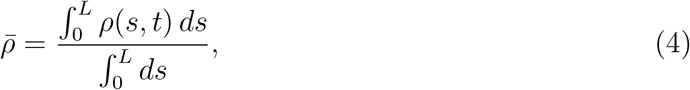

where *L* = *a* + *b* + *ℓ* is the total length of the membrane region in which regulators are distributed (see Fig. 1B, right), and *a*, *b*, and *ℓ* are the respective apical, basal, and lateral lengths of the cell (see Fig. 1A).

### B. The dependence of polarity formation on cell geometry

The epithelial polarity regulators either reside in the cytosol or are associated with the membrane surface. As indicated in Eq. (3), for a given polarity regulator, the probability of residing in the cytosol versus associating with the membrane depends on the surface-to-volume ratio *ψ* of the cell, which is calculated as follows:

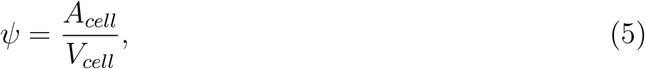

where *V_cell_* is the cell volume, and *A_cell_* = *A_a_* + *A_b_* + *A_ℓ_* is the total surface area of a cell with *A_a_* the apical area, *A_b_* the basal area, and *A_ℓ_* the lateral area. Moreover, the half cell perimeter *L* also affects epithelial membrane polarization as it determines the average membrane concentration of the polarity regulators, as revealed by Eq. (4).

### C. The mechanical module for cell shape changes in three dimensions

To simulate 3D epithelial shape formation, we developed a vertex-based mechanical model for a monolayered epithelial tissue consisting of *N* cells. In the vertex model, the full 3D shape of a cell is described by a hexagonal prism with edges and vertices [1] (Fig. 1A). Forces acting on the vertices are derived from the derivative of the potential function *U*, which describes the mechanical properties of the entire tissue. Such forces are balanced by the surrounding viscous drag in an environment of low Reynolds numbers. Thus, the equation of motion for cell shape changes reads,

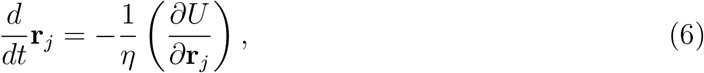

where **r**_*j*_ is the position coordinate of vertex *j*, and *η* is the friction coefficient. A force-balanced tissue morphology is achieved when *U* reaches a minimum.

The potential function that describes the mechanics of the entire tissue takes into account energy contribution from all cell surfaces and cell volumes. Specifically, we considered energy resulting from the surface tensions on the basal and lateral sides of the cells, the volume elasticity that maintains the cell volume during cell shape changes, and the elastic energy of the apical surface given that cytoskeletons (e.g., actomyosin and microtubule networks) are enriched on the apical sides of the cells [27]. The potential function is thus formulated as follows,

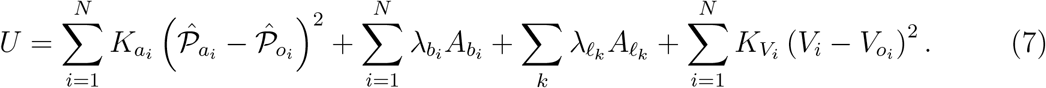

The first term describes the apical elasticity with elastic modulus 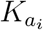, apical perimeter 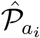, and preferred perimeter 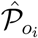 for the *i*-th cell. The second and third terms describe the tension along the basal and lateral surfaces, respectively, with surface tension 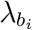 and area 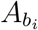 for the basal surface, while surface tension 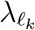 and area 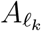 for the lateral surface with an index *k* that runs across all lateral cell surfaces. The fourth term defines the volume elasticity with elastic modulus 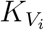, cell volume *V_i_*, and preferred volume 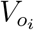 for the *i*-th cell. The volume elasticity likely arises from the hydrostatic pressure of the cytoplasm and the intracellular cytoskeletons that resist changes of the cell volume. Note that perimeter 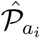, area 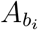 and 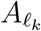, as well as volume *V_i_* are calculated from the vertex positions, whose dynamics follow Eq. (6).

By using qualitatively distinct mechanical terms, we took into account that the apical and basal surfaces of a polarized epithelial cell possess distinct properties. In particular, in addition to using a tension term for the apical perimeter as was used previously [15], we further considered the elastic properties of the cytoskeletons along the apical perimeter. Such apical elasticity helps to stabilize cell shape under large deformation.

### D. Polarity-dependent modulation of cell mechanics as a result of apical domain homeostasis

Motivated by the observation that changes in epithelial cell polarity initiate tissue folding during *Drosophila* gastrula [40], we devised a mechanism for polarity-dependent modulation of cell mechanical properties. Previous work on dorsal fold formation found that while down-regulation of the polarity regulator Par-1 initially triggers the expansion of the aPKC domain, as defined by the region apical to the position of Bazooka localization, on longer timescales during cell shortening that initiates fold formation, the length of the aPKC domain remains relatively constant, revealing homeostatic maintenance in the apical domain [38]. It was thus proposed that as the Bazooka position continues to shift towards the basal side of the cells due to Par-1 downregulation, the mechanism that ensures a constant aPKC domain size during the basal shifts of Bazooka enables the reduction of cell height [38]. Because the mechanism underlying the aPKC domain homeostasis remains unclear, we modeled the mechanistic link between cell polarity and mechanics using a phenomenological approach. Specifically, we hypothesized a functional term that defines the polarity-dependent modulation of cell mechanics based on the homeostatic relaxation of the aPKC domain, as follows:

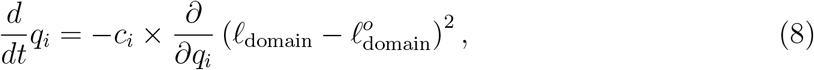

where 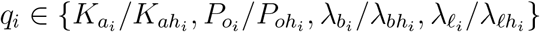 is the normalized mechanics of cell *i* with respect to the initial values 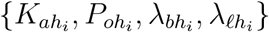, and *c_i_* is a parameter which quantifies the rate of change of cell mechanics *q_i_* induced by polarity remodeling. *ℓ*_domain_ is the aPKC domain size, calculated numerically as the length from the apical surface to the position of maximum Bazooka density (see Fig. 1B, right), and 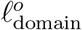 is the prescribed homeostatic domain size. Equation (8) thus phenomenologically describes cell mechanical modulation in a manner that accompanies the reduction in size variation of the homeostatic apical domain.

In sum, the polarization of epithelial cell membrane is described by Eq. (1), where the polarity formation depends on the whole cell shape, as indicated by Eq. (3). The changes in cell shape follow Eq. (6), while the forces that deform the cells are in turn regulated by the cell polarity via Eq. (8).

## III. RESULTS

### A three-element network model recapitulates key features of epithelial apical-basal polarization in the *Drosophila* cellular blastoderm

The *Drosophila* cellular blastoderm forms its primary epithelium via cellularization. During cellularization, the microtubule-dependent, minus end-directed transport of Bazooka, and its eventual accumulation on the apical side of the cell, function as the key symmetry breaking event that polarizes the cells [18]. To simulate the observed apical accumulation of Bazooka, we used the Heaviside step function for the initial distribution of *B*, while keeping *A* and *P* uniform along the apical–basal axis (Fig. 2A, dashed lines). We then numerically solved the spatio-temporal distribution of polarity regulators along the cell perimeter with periodic boundary conditions according to Eq. (1) and parameter values given by Table I. As shown in Fig. 2A, the distribution of *A*, *B* and *P* evolves toward a steady-state polarized profile, indicating that the *A*–*B*–*P* network recapitulates epithelial polarization in the early *Drosophila* embryo. The simulated steady-state profile resembles that observed in the early *Drosophila* embryo, in the sense that while aPKC (*A*) and Par-1 (*P*) form the polarized apical and basal-lateral domains, respectively, Bazooka (*B*) is localized as a sharp peak between these two domains.

**TABLE I.**
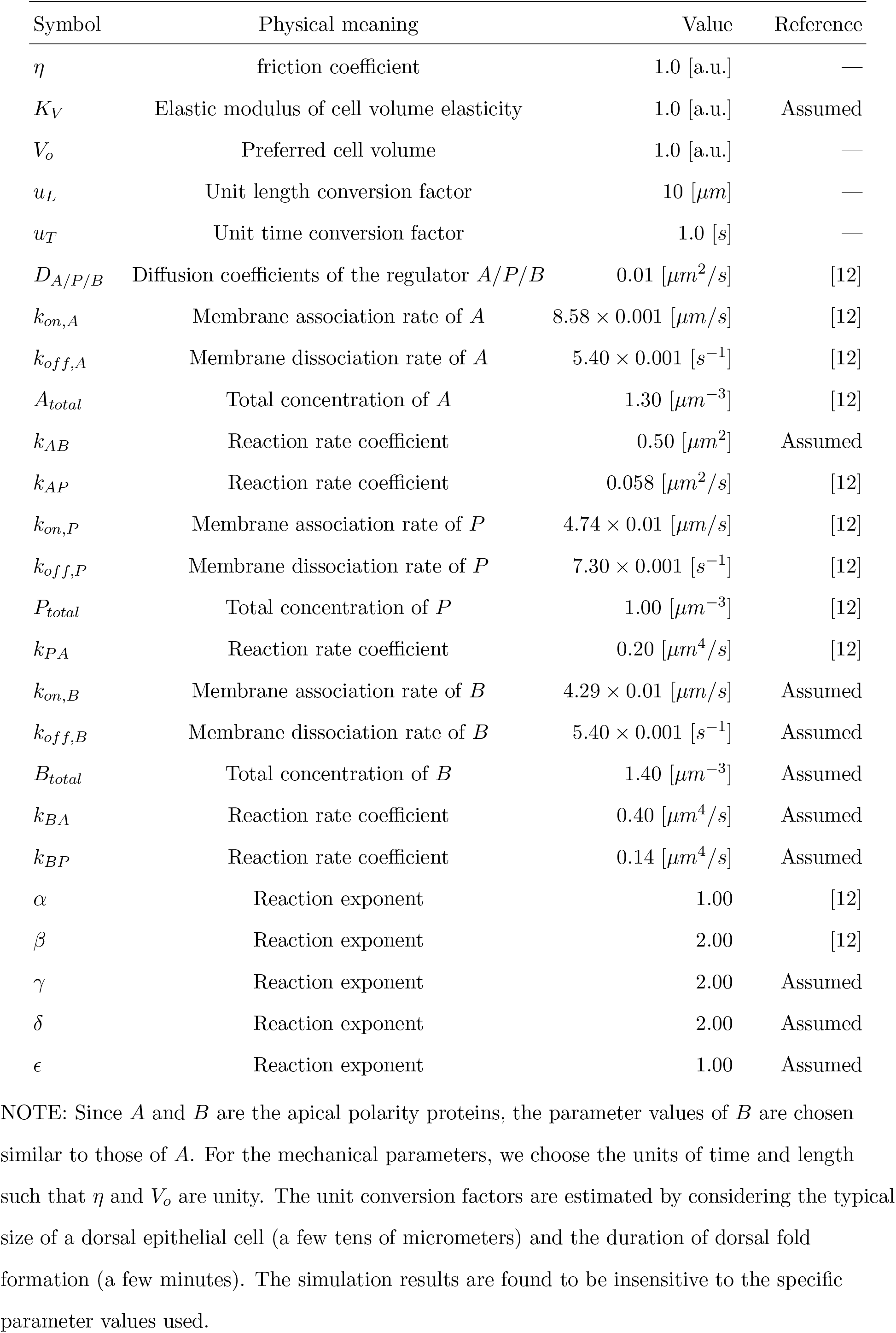
Parameters used for the mechano–biochemical simulations

| Symbol | Physical meaning | Value | Reference |
| --- | --- | --- | --- |
| $\eta$ | friction coefficient | 1.0 [a.u.] | — |
| $K_V$ | Elastic modulus of cell volume elasticity | 1.0 [a.u.] | Assumed |
| $V_o$ | Preferred cell volume | 1.0 [a.u.] | — |
| $u_L$ | Unit length conversion factor | 10 [$\mu m$] | — |
| $u_T$ | Unit time conversion factor | 1.0 [s] | — |
| $D_{A/P/B}$ | Diffusion coefficients of the regulator $A/P/B$ | 0.01 [$\mu m^2/s$] | [12] |
| $k_{on,A}$ | Membrane association rate of $A$ | $8.58 \times 0.001$ [$\mu m/s$] | [12] |
| $k_{off,A}$ | Membrane dissociation rate of $A$ | $5.40 \times 0.001$ [$s^{-1}$] | [12] |
| $A_{total}$ | Total concentration of $A$ | 1.30 [$\mu m^{-3}$] | [12] |
| $k_{AB}$ | Reaction rate coefficient | 0.50 [$\mu m^2$] | Assumed |
| $k_{AP}$ | Reaction rate coefficient | 0.058 [$\mu m^2/s$] | [12] |
| $k_{on,P}$ | Membrane association rate of $P$ | $4.74 \times 0.01$ [$\mu m/s$] | [12] |
| $k_{off,P}$ | Membrane dissociation rate of $P$ | $7.30 \times 0.001$ [$s^{-1}$] | [12] |
| $P_{total}$ | Total concentration of $P$ | 1.00 [$\mu m^{-3}$] | [12] |
| $k_{PA}$ | Reaction rate coefficient | 0.20 [$\mu m^4/s$] | [12] |
| $k_{on,B}$ | Membrane association rate of $B$ | $4.29 \times 0.01$ [$\mu m/s$] | Assumed |
| $k_{off,B}$ | Membrane dissociation rate of $B$ | $5.40 \times 0.001$ [$s^{-1}$] | Assumed |
| $B_{total}$ | Total concentration of $B$ | 1.40 [$\mu m^{-3}$] | Assumed |
| $k_{BA}$ | Reaction rate coefficient | 0.40 [$\mu m^4/s$] | Assumed |
| $k_{BP}$ | Reaction rate coefficient | 0.14 [$\mu m^4/s$] | Assumed |
| $\alpha$ | Reaction exponent | 1.00 | [12] |
| $\beta$ | Reaction exponent | 2.00 | [12] |
| $\gamma$ | Reaction exponent | 2.00 | Assumed |
| $\delta$ | Reaction exponent | 2.00 | Assumed |
| $\epsilon$ | Reaction exponent | 1.00 | Assumed |

NOTE: Since $A$ and $B$ are the apical polarity proteins, the parameter values of $B$ are chosen similar to those of $A$ . For the mechanical parameters, we choose the units of time and length such that $\eta$ and $V_o$ are unity. The unit conversion factors are estimated by considering the typical size of a dorsal epithelial cell (a few tens of micrometers) and the duration of dorsal fold formation (a few minutes). The simulation results are found to be insensitive to the specific parameter values used.

**FIG. 2.**
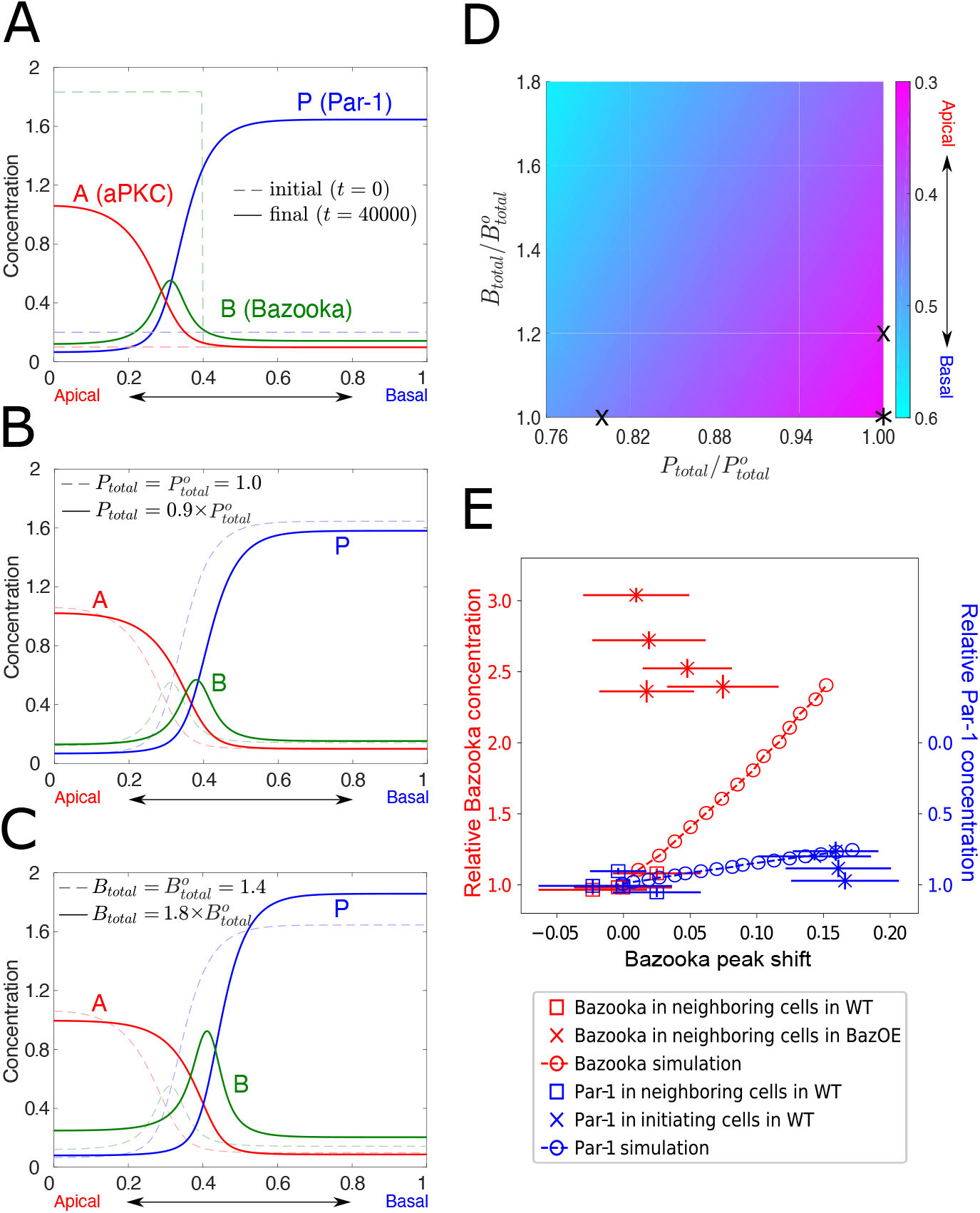
The ”*A*–*B*–*P* network” model recapitulates the spatial profile of apical–basal polarity formation and remodeling in the *Drosophila* cellular blastoderm. (A) The initial and final steady-state distributions of polarity regulators in the model. The membrane location is scaled by *L* and hereafter. (B, C) Basal shifts of the Bazooka peak position at the steady state (solid line) following a decrease in *P_total_* (B) or an increase in *B_total_* (C) as compared to the steady-state profile (dashed line) in (A). (D) Color plot of the Bazooka peak positions as a function of fold change of *P_total_* and *B_total_*. The crosses are for the comparison between Bazooka peak positions with a 20% perturbation of *P_total_* decrease and *B_total_* increase. The asterisk marks the unperturbed control. (E) Bazooka peak position shift in response to Par-1 down-regulation (blue, y-axis right) or Bazooka overexpression (red, y-axis left) in experiments (squares and crosses with error bars) and simulations (circles with a dashed line). Each experimental datapoint represents the mean measurement from one embryo. Error bars are standard deviations of sample sizes *n* = 20–30 for the initiating cells or *n* = 70–100 for the neighboring cells. See Materials and methods for details on experimental measurements.

We next asked whether the *A*–*B*–*P* network model recapitulates features of polarity remodeling previously observed in the experiments. On the dorsal side of cellularizing embryo, down-regulation of Par-1 in a specific population of cells, called the initiating cells, results in basal shifts of the positions of Bazooka and adherens junctions [40]. To simulate this, we introduced a 10% reduction in the Par-1 total concentration to the above steady-state polarity profile in Fig. 2A, approximating the observed decrease of Par-1 in the initiating cells prior to the end of cellularization [40]. Figure 2B shows that the peak position of *B* exhibits a basal shift of approximately 10% cell height, comparable to the experimental observations [40]. Given that Bazooka co-localizes with adherens junctions in the early *Drosophila* embryo [17, 18], the peak position of *B* corresponds to the junctional position. This result thus supports the hypothesis that basal shifts of adherens junctions could result from biochemical remodeling of polarity regulators, i.e. the down-regulation of Par-1 [40].

Previous work also showed that overexpression of Bazooka is sufficient to induce basal shifts of junctions that cause the formation of ectopic dorsal folds [40]. Motivated by this, we increased the total concentration of *B* in the steady-state profile shown in Fig. 2A, and found that the peak position of *B* shifts basally by 10% when *B* concentration is increased by 80% (Fig. 2C), consistent with experimental observations reported previously [40]. The result also raises the question of whether the system exhibits quantitatively different response to Par-1 down-regulation and Bazooka overexpression. To address this question, we explored a wide range of *P_total_* and *B_total_* changes. For the same magnitude of variation, we observed a greater shift in the peak position of *B* in response to changes in *P_total_* than *B_total_* (Fig. 2D; see the crosses for comparison of a 20% variation). This result predicts that the position of adherens junctions is more sensitive to changes in the concentration of Par-1 than Bazooka.

To test this theoretical prediction, we re-analyzed a previously published dataset on the extent of junctional shifts in response to Par-1 and Bazooka changes [40]. For Par-1 down-regulation, we compared wild-type (WT) neighboring cells (i.e., high Par-1) with WT initiating cells (i.e., low Par-1), whereas for Bazooka overexpression (BazOE), we compared WT neighboring cells (i.e., low Bazooka) with BazOE neighboring cells (i.e., high Bazooka) (see Materials and methods for details). Figure 2E shows that greater basal shifts of junctions are observed with Par-1 down-regulation than with Bazooka overexpression. Such experimental results are in good agreement qualitatively with the theoretical predictions (pointed-dashed lines), in the sense that changes in Par-1 concentration have a stronger impact on the positioning of junctions than changes in Bazooka concentration. Notably, the effect of junctional shifts following Par-1 downregulation is found to be quantitatively comparable between the theory and experiments. We observed, however, greater junctional shifts in our theory, as compared with the experimental observation, following Bazooka overexpression. This may result from experimental measurements of Bazooka overestimating the effective concentration of Bazooka involved in polarity formation. Alternatively, the experimental overexpression of Bazooka may have exceeded the physiological range of Bazooka levels that induces additional regulatory mechanisms to saturate the system.

### Cell geometry exerts two distinct effects – scaling and remodeling – on the polarity profile

Epithelial cells could increase or decrease in volume, lengthen or flatten as the aspect ratio changes, or undergo anisotropic deformation due to the contraction or expansion of a specific membrane surface. To understand the impact of geometric changes, we considered the effect of two specific cell features, surface-to-volume ratio *ψ*, Eq. (5), and half cell perimeter *L* (see the thick black line in Fig. 1A and the right panel of Fig. 1B), on polarity distribution under the assumption that the total concentrations of polarity proteins are constant. As shown in Fig. 3A, an increase in *ψ*, which happens when a cell deforms from a cuboidal to a columnar shape, induces a basal shift of polarity. In contrast, although an increment in *L* also induces basal shifts of polarity (Fig. S1), the peak position of *B* scales with *L* (Fig. 3B).

**FIG. 3.**
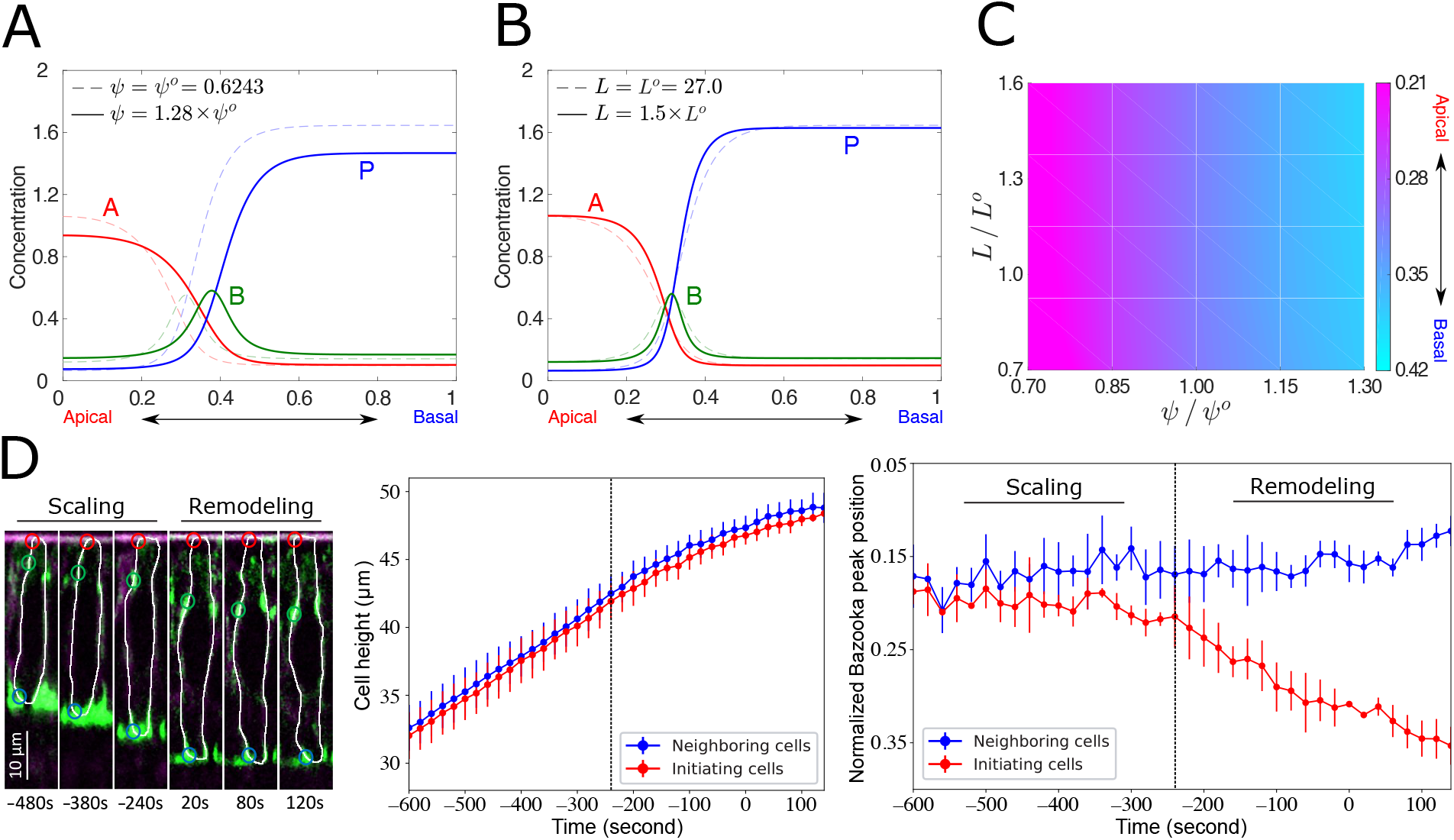
Cell geometric features affect membrane distribution of the polarity regulators. (A, B) The Bazooka peak position at the steady state (solid line) shifts basally following an increase in *ψ* (A), whereas remains unchanged when *L* is increased (B) as compared to the control (dashed line). (C) Color plot of the Bazooka peak positions as a function of fold change of *ψ* and *L*. (D) (Left) Time-lapse images of a representative initiating cell outlined in white in the embryo expressing E-Cadherin-3xGFP, Myosin-GFP and Gap43-mCherry. The red, green, and blue circles label the apical, junctional, and basal points, respectively, for quantification of the cell height and the Bazooka peak positions. (Middle) Time series of cell height averaged over the initiating (red) or neighboring cells (blue). (Right) Time series of Bazooka peak positions for both the initiating (red) and the neighboring cells (blue). In the middle and right panels, the vertical dash lines are plotted manually, serving as visual guides for the boundary between the scaling and remodeling phases. The bars of each data point are the standard deviation of the datasets derived from three embryos. See details of experimental setup and quantification method in Materials and methods.

Our theory thus predicts two distinct types of basal polarity shifts: (1) scaling, in which the relative junctional position remains unchanged during polarity redistribution (e.g., see Fig. 3B), and (2) remodeling, in which shifts of the relative junctional position alter the proportion of the apical to basal-lateral membrane domains (e.g., see Fig. 2, B and C, and Fig. 3A). Morphogenesis of dorsal epithelial cells during early *Drosophila* development could be used to test these predictions. Prior to dorsal fold initiation, all dorsal epithelial cells lengthen as cellularization proceeds, while maintaining the width, thereby increasing *L* (see Fig. 3D, left and middle), with uniform levels of Par-1 across cell populations. Approximately 7 minutes prior to the end of cellularization, Par-1 levels begin to decrease in the initiating cells, likely due to an induced downregulation, causing a decrease of *P_total_*, but the Par-1 level remains constant in the neighboring cells [40]. Our model thus predicts that both initiating and neighboring cells exhibit the scaling effect during early to mid cellularization. Towards the end of cellularization, however, as active downregulation of Par-1 begins in the initiating cells, the scaling phase is expected to end, whereas the junctional position in the neighboring cells should continue to scale with the cell height. In accordance with this, quantitative live imaging of *Drosophila* dorsal epithelial cells (see details in Materials and methods) showed that the junctional position scales with cell height in both the initiating and neighboring cells as cellularization proceeds (Fig. 3D, right). As cellularization approaches the end, the junctional positions in the initiating cells begin to exhibit greater basal shifts, deviating from the scaled position, while those in the neighboring cells remain scaled with *L*. Thus, our experimental data validate the theoretical prediction on the scaling effect. Note also that for the initiating cells, the initial basal shifts of junctions have been shown to occur without cell shortening [40]. Since *ψ* is expected to be near constant during these initial basal shifts, these shifts must result from biochemically induced downregulation of Par-1, but not the *ψ*–dependent remodeling effect. Following such biochemically initiated basal shifts, the initiating cells begin to shorten into a frustum shape, during which it is possible that the increased *ψ* resulting from the shortening (as will be shown later in Fig. 4A) further enhances the basal shifts.

**FIG. 4.**
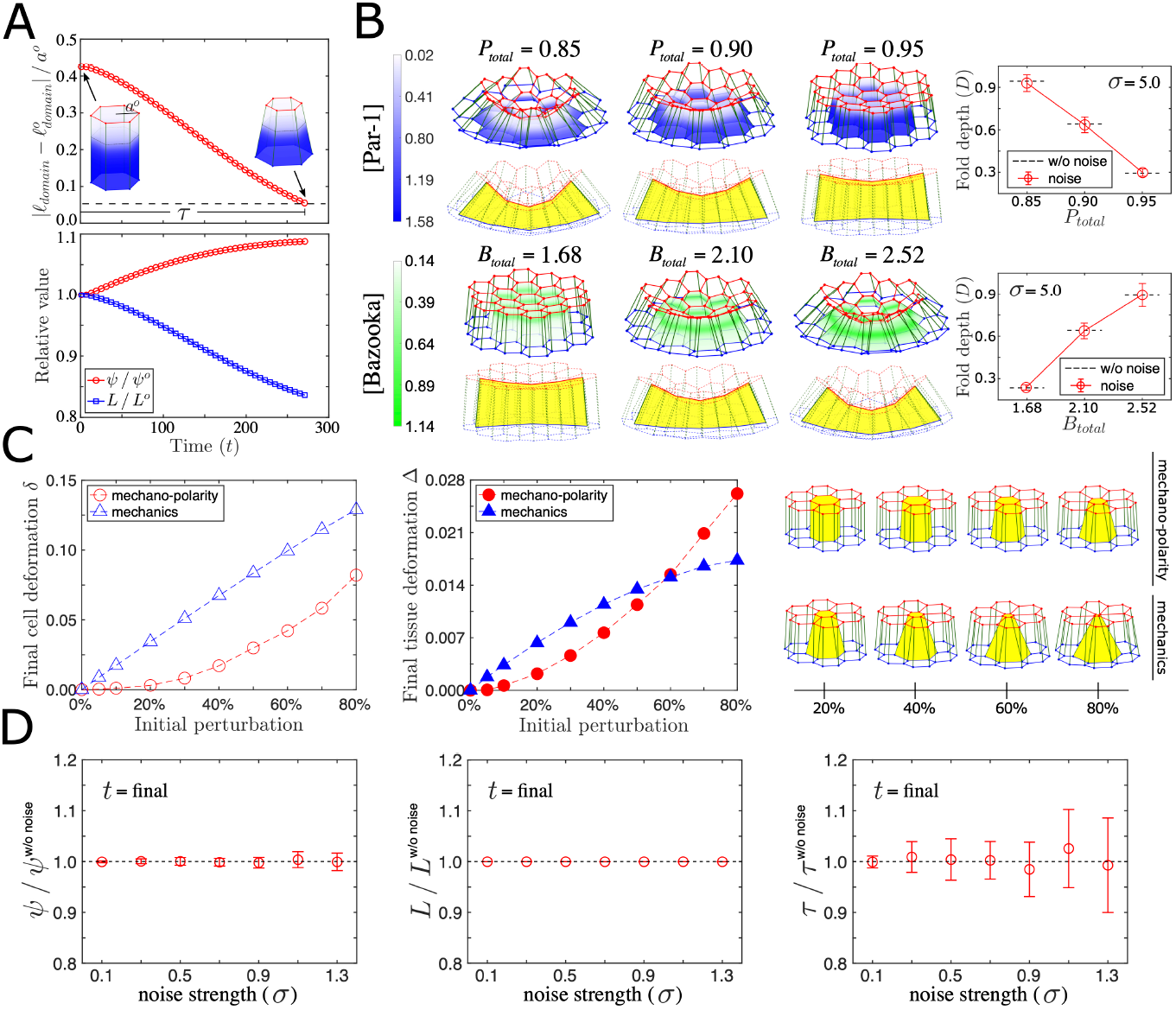
Mechano–polarity feedback induces spontaneous and robust cell shape change and tissue deformation. (A) The deviation of apical domain (*ℓ*_domain_) from its homeostatic 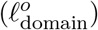 size following a decrease (*P_total_* = 0.9) of total concentration of *P* plotted as a function of time (top) with the corresponding time evolution of *ψ* and *L* normalized by their initial values (bottom). Horizontal dashed line depicts the threshold of the apical homeostasis, and *τ* denotes the time required to reach the threshold. Insets: the initial and homeostatic 3D cell shapes with the graded color showing *P* concentration (see color code in (B)) and the dotted lines showing the peak positions of *B*. (B) 3D folded epithelia together with a cross-sectional view (left columns) and quantitation of steady-state fold depths (far right column) following localized decrease of *P_total_* (top row) or increase of *B_total_* (bottom row). Color gradients indicate the steady-state profile of *P* (blue) and *B* (green) concentrations in the cells that are initially subject to the polarity change. The fold depths without noise (dashed lines) were plotted together with the mean fold depths of 10 different simulations with noise strength *σ* = 5.0 (circles with error bars). The error bars are standard deviations depicting the variabilities of fold depths. (C) Final cell deformation *δ* (left) and final tissue deformation Δ (center) plotted against the initial perturbation on the apical mechanics 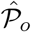 with and without mechano– polarity feedback. Final steady-state tissue morphology for different initial mechanical perturbations (right). See Materials and methods for details of the analysis. (D) Cell shape features, *ψ*and *L* (left and middle columns), and the relaxation time, *τ* (right column), plotted against *σ*. The quantities *ψ*, *L* and *τ* are normalized by those without noise, and averaged over 10 different simulations with a given noise strength (circles with error bars). The error bars are standard deviations of the simulation results. 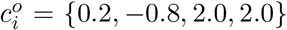 for (B), 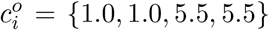 for (C), 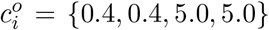 and for (D), and others given in Table I are used in the simulations.

### A feedback loop between cell mechanics and polarity leads to spontaneous tissue fold formation

So far we have shown that changes in epithelial cell shape alter the distribution of polarity regulators along the cell membrane, and as reported in Appendix A, cell shapes themselves depend on the mechanical properties of the membrane surfaces. The apical polarity domain homeostasis observed in the *Drosophila* dorsal fold system ([38]; see also the texts associated with Eg. (8) in the Mathematical Modeling section) suggests that polarity redistribution may feed back on the modulation of cell surface mechanics, which can further change the cell shape. Specifically, for the *Drosophila* dorsal epithelial cells, polarity distribution could be influenced by the geometric features of the cell shape, while the cell shape itself could be controlled by cell mechanics that in turn depends on cell polarity. Thus, cell mechanics, cell shape and cell polarity could potentially be linked in a feedback loop.

To study how this mechano–polarity feedback loop controls epithelial cell shape, we numerically computed both polarity distribution and cell shape based on Eqs. (1) and (6) with their coupling terms given by Eqs. (5) and (8). Modulation was applied to cells that initially have a stable shape and steady-state polarity distribution. To simulate the condition of Par-1 down-regulation (or Bazooka overexpression), we decreased (or increased) total concentration of *P* (or *B*). This induces an initial expansion of the *A* domain, which subsequently shrinks toward its homeostatic size (Fig. 4A, top). During such homeostatic relaxation, surface-to-volume ratio *ψ* increases, while half cell perimeter *L* decreases (Fig. 4A, bottom). Ultimately, cells transform from a tall columnar shape to a short frustum shape, resembling the shape changes that the initiating cells exhibit during *Drosophila* dorsal fold initiation [40]. Since cells in the *Drosophila* dorsal fold system initially have columnar shapes, during mechano–polarity feedback coupling, homeostatic relaxation of the apical domain can be most effectively achieved by the alteration of cell height.

We next examined the tissue-scale morphological changes following alterations of cell polarity in this mechano–polarity feedback system. Figure 4B shows that both a decrease in the total concentration of Par-1 (*P_total_*) and an increase in that of Bazooka (*B_total_*) within a localized tissue area induce out-of-plane deformation, recapitulating both normal and ectopic dorsal fold formation [40]. Thus, a local trigger of polarity remodeling can induce spontaneous epithelial folding via mechano–polarity feedback without external forces. These results provide a theoretical foundation for the previously proposed hypothesis that dorsal fold formation is initiated cell-autonomously, whereas external forces from the surrounding tissues are dispensable [40].

### Mechano–polarity feedback endows cells with tolerance to mechanical/biochemical perturbations

To further explore the properties of this mechano–polarity feedback loop, we perturbed the apical mechanics of an initially stable cell (highlighted in yellow in Fig. 4C) by modulating 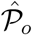 with a fixed change of *K_a_*, and assayed the corresponding shape deformation both at the cell and tissue scales. Remarkably, when the strength of the initial perturbation is low, only slight cell and tissue deformation are observed, whereas the deformation response increases dramatically as the perturbation strength increases (Fig. 4C, red pointed-dashed lines on the left and middle panels). In contrast, the response of cell and tissue deformation to the mechanical perturbation lacks such a near refractory regime at low perturbation strengths when the system does not contain mechano–polarity feedback (see the blue pointed-dashed lines on the left and middle panels of Fig. 4C for comparison).

This result suggests that the mechano–polarity feedback loop endows cells with a regime within which small mechanical fluctuations, presumably arising stochastically, lead to minuscule, non-consequential morphological changes, whereas active, instructive mechanical alterations exceeding a certain level are required to induce substantial morphological changes, implying that the mechano–chemical coupling could generate a threshold-like response, such that only perturbations above the threshold value can induce sufficiently large deformation. Thus, a system with mechano–polarity coupling could be more resistant to the noise-induced ectopic initiation of morphogenesis. Note that a sharp cell/tissue deformation response was also observed following the perturbation of Par-1 total concentration (Fig. S2). This result suggests that low-amplitude fluctuations of Par-1 concentration at the steady-state do not lead to epithelial fold initiation, whereas genetically programmed down-regulation of Par-1 that provides a sufficiently strong perturbation would be required to trigger epithelial folding.

Notably, as revealed in Fig. 4C (left and middle panels), whereas cell shape change is smaller with mechano–polarity coupling (red line) than without (blue line), for sufficiently strong perturbations, the coupled system generates greater tissue deformation than the system that lacks coupling. This result can be understood as the consequence of the subsequent modulation of basal–lateral mechanics following the initial perturbation of apical mechanics due to mechano–polarity coupling, given that apical modulation induces only a shallow tissue indentation, whereas basal–lateral modulation generates a deeper fold (Fig. S3A). Thus, mechano–polarity coupling allows modulation of mechanics at membrane surfaces that are not the primary target of the initial mechanical stimuli and may promote the efficiency of tissue deformation.

### Tissue deformation driven by mechano–polarity feedback is robust against stochasticity

Lastly, we sought to determine whether mechanical fluctuations render the self-organized epithelial deformation stochastic as biological systems are intrinsically noisy. We added noise to the mechano–polarity coupling term *c_i_* to simulate fluctuations in the rate of conversion from the polarity signal to the mechanical conditions at the cell membrane. The noise is given as follows:

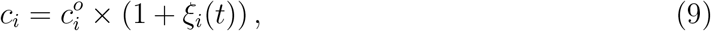

where 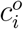 is the deterministic portion of the polarity-to-mechanics conversion rate, while *ξ_i_*(*t*) is the Gaussian white noise that has a zero mean and a standard deviation *σ*. The value of *σ* defines the extent of stochasticity in how cell mechanics *q_i_* decode the polarity signal (see Eq. (8)).

To evaluate robustness at the cellular level, we quantified the geometric features, *ψ* and *L*, of the final cell shape, and the time, *τ*, it takes to achieve apical domain homeostasis. We observed that both *ψ* and *L* are nearly identical across a wide range of noise strengths *σ* (Fig. 4D, left and middle), although the variability of relaxation time *τ* increases with stronger noise (Fig. 4D, right). At the tissue level, we also observed low levels of variability in the epithelial fold depths (Fig. 4B, right). These data indicate that mechano-polarity coupling ensures robust cell shape change and tissue folding despite noise.

## VI. DISCUSSION

This work proposes a theoretical model for dorsal fold formation in the *Drosophila* embryo, an epithelial folding event that is induced by changes in epithelial cell polarity. Our model recapitulates key features of polarity remodeling and links changes in cell polarity to mechanical modulation by phenomenologically mimicking the experimentally observed apical domain homeostasis. Such mechanical modulation induces cell shape changes, altering system geometry on which the reaction-diffusion process of the polarity network depends, thereby forming a feedback loop of mechano–polarity coupling. Simulation with this feedback model indicates that genetically programmed Par-1 down-regulation can trigger spontaneous cell shape changes, leading to self-organized tissue fold formation. Such an intracellular mechano–polarity crosstalk prevents epithelial tissues from undergoing shape changes under small fluctuations in cell mechanics and polarity, which could occur owing to the intrinsic noise that may arise during development. The intrinsic property of robustness against stochastic fluctuations can thus ensure that the initiation of autonomous tissue deformation is spatio-temporally precise and tightly controlled.

The current model contains two key hypotheses for which mechanistic insights remain to be explored. First, we hypothesized that surface-to-volume ratio *ψ* affects the balance between the cytosolic and membrane pools of the polarity regulators. Previous studies have reported that membrane localization of the PAR proteins depends on their association with the specific lipid moieties in the cell membrane. For instance, Par-1 binds to phosphatidylserine [29], whereas Bazooka mainly associates with phosphatidylinositol [13, 26]. Assuming that in the membrane region in which the polarity proteins localize, the percentage presence for each specific type of lipid is near constant, the number of these ”membrane receptors” for a given polarity protein would likely scale proportionally with the membrane surface area. The constant membrane lipid composition could thus be the potential mechanism that underlies *ψ*-dependent cytosol–membrane exchange of the polarity proteins. Note that variation of *ψ* has also been proposed to create biochemically distinct states (e.g., with regard to the phosphorylation status of a protein) within a cell, which influences cell division timing or the scaling of internal organelles to the overall cell sizes [19, 28]. We speculate that alterations of surface-to-volume ratio can be a common mechanism that modulates intracellular reactions, thereby regulating cellular behaviors.

The second hypothesis is that aPKC domain homeostasis underlies mechanical modulation in response to polarity change. As mentioned in the Mathematical Modeling section, a recent study on the initiation of dorsal fold formation reveals that the apical domain of the initiating cells maintains a relatively constant size as the cells undergo shortening to induce fold formation [38]. The study also reveals potential connections between the polarity proteins and the mechanical elements involved in the homeostasis. Specifically, the microtubule minus-end binding protein Patronin was shown to become associated with the cell cortex underneath the apical surface in an aPKC-dependent manner. The cortex-associated Patronin organizes an apical microtubule network that appears to undergo rapid turnover, which depends on Patronin-dependent recruitment of the microtubule-severing enzyme Katanin that likely destabilizes the microtubules. Given that Patronin distribution in the apical domain responds to downregulation of Par-1 during dorsal fold initiation, and the forces that the apical microtubule network exerts on the apical cortex may scale with the cortical density of Patronin, it was proposed that this rapid turnover, coupled with the polarity-dependent cortical localization of Patronin and the apical microtubule network that associates with it, underlies the apical size homeostasis [38]. Patronin could thus be at the nexus of this mechano–polarity coupling in the dorsal fold system. It is worth noting that in the Patronin RNAi embryos the shape of the dorsal epithelial cells become extremely irregular [38]. This observation is consistent with the prediction of our model that without mechano–polarity coupling the cell shapes are more sensitive to fluctuations in mechanics and cell polarity. We propose that the reduced Patronin activity impairs mechano–polarity coupling such that cells in the epithelium become more prone to the ambient noise in the system.

Although we constructed the current model by closely following properties specifically linked to the *Drosophila* dorsal fold system, it is likely that the systems-level characteristics of the mechano–biochemical crosstalk, including the autonomy of tissue deformation and the robustness against noise, transcend context-specific biochemical mechanisms of cell polarity and mechanical regulation of cell shapes. In support of this, self-organized behaviors orchestrating cell shapes have been observed in previous models that consider bidirectional feedback between cell mechanics and tissue-scale polarity cues [35] or morphogen patterns [4, 6, 31]. Moreover, it has been proposed that feedback between intracellular mechanics and extracellular morphogen concentrations leads to robust formation of morphogen concentration profile [4, 6], thereby ensuring proper tissue architecture. We envision that self-organization and robustness are general properties of mechano–chemical feedback control during tissue morphogenesis and remodeling. Our modeling framework could potentially serve as a generalizable platform that can be modified to model mechanical–biochemical coupling systems whose mechanistic details are distinct from the dorsal fold system.

The vertex-based modeling represents a powerful framework for simulating tissue morphogenesis in various biological contexts [1, 9]. The vertex models have recently been developed to further take into account the regulatory mechanisms of cell mechanics. In contrast to previous 3D models that consider regulation of cell shape changes by a global, tissue-scale signaling pattern [6, 31], cell deformation in our model is regulated by a local signal resulting from polarity pattern formation within a cell, which is in turn affected by cell deformation. Thus, our model provides insights into regulation of cell mechanics that arises at the cellular scale, while adjustable to the resultant change of cell shape, and reveals novel tissue properties emerging from such intracellular crosstalk.

## MATERIALS AND METHODS

### A. Quantitation of junctional positions in the dorsal epithelium of fixed wild-type or Bazooka overexpression embryos

Previously published datasets [40] were re-analyzed for the comparison between the effects of Par-1 down-regulation and Bazooka overexpression on the position of adherens junctions (represented as the Bazooka peak position). Par-1 and Bazooka concentration was quantified as previously described [40]. In brief, wild-type embryos and embryos overexpressing Bazooka were fixed, stained with Par-1, Bazooka and Neurotactin (for cell membrane visualization) antibodies, and imaged in 3D on a Leica SP5 scanning confocal system with a 63x glycerol immersion objective. The resultant images were processed, segmented and quantified using the EDGE4D software [23]. Bazooka concentration was quantified as the per voxel intensity within the segmented Bazooka junctional localization. Par-1 concentration was the per voxel intensity within a two-voxel distance from the 3D reconstructed membrane surface mesh in the region basal to the Bazooka junctions. The relative concentration was computed by normalizing the concentration of Par-1 or Bazooka by the mean concentration of Par-1 or Bazooka in the neighboring cells of all wild-type embryos. The Bazooka peak position was normalized by the total cell height. The shift of Bazooka peak was measured as the normalized position relative to its mean value in the neighboring cells of all wild-type embryos.

Junctional shifts induced by Par-1 down-regulation in the wild-type embryos were inferred by a comparison between normalized Bazooka peak positions in the initiating cells of the posterior fold and those in the neighboring cells anterior to the posterior folds. Junctional shifts induced by Bazooka overexpression were estimated by a comparison between wild-type and Bazooka overexpression embryos of the normalized Bazooka peak positions in the neighboring cells.

### B. Quantitative analysis of time evolution of junctional positions during *Drosophila* dorsal fold formation

Embryos expressing E-Cadherin-3xGFP [33], Myosin-GFP [34], and Gap43-mCherry [22] were imaged on an Olympus Fluoview FVMPE-RS multi-photon scanning system with a 20X water immersion objective. The resultant images were processed, segmented and quantified using a custom-made ImageJ macro. Because of the slender columnar cell shape, junctional position (i.e., Bazooka peak position) was estimated as the mean vertical distance between the pixels with local maxima of E-Cadherin-3xGFP intensity (i.e., the green circles in Fig. 3D), given that Bazooka and E-Cadherin positions are identical [40], and the basal margin of the segmented autofluorescent intensity of the vitelline membrane (i.e., the red circles in Fig. 3D), defined as the cell apices. Cell height, which approximates *L* in the theory, was estimated as the mean vertical distance between the bases of the cells, defined as the basal margin of the segmented Myosin-GFP intensity (i.e., the blue circles in Fig. 3D), and the cell apices. Gap43-mCherry allows for visualization of cell membrane. The region centered on the posterior fold spanning approximately 2–3 cells was used as the ROI representing the initiating cells, whereas the neighboring cells were represented by a region of 2–3 cells located 4–5 cells anterior to the initiating cells of the posterior fold. Quantitation was based on the time series of a single image of XZ line scan (i.e., the sagittal plane), followed by temporally aligning the resultant datasets (n = 3) at time zero defined as the end of cellularization in each respective embryo. The end of cellularization was estimated as the time at which the cell height of those in the region anterior to the cephalic furrow reaches a maximum.

### C. Quantification of the perturbation–deformation relationship

In Fig. 4C and Fig. S2, we studied cell and tissue deformations in response to mechanical or biochemical perturbations, respectively. For an initially balanced epithelium, either the apical mechanics, 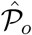 and *K_a_*, or the total Par-1 concentration, *P_total_*, was modulated in the central cell (highlighted in yellow in Fig. 4C and Fig. S2) at time *t* = 0, where *K_a_* = 9 × *K_ac_* and *K_ac_* is the original unperturbed value of the apical elastic modulus. The initial perturbation strength of 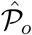 and *P_total_* were quantified as 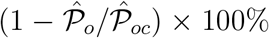 and 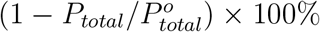, respectively, with 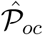 and 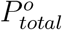 being their unperturbed values. The final cell deformation was quantified as the changes of each cell surface relative to the unperturbed control, calculated as 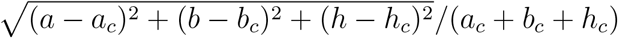, where *a_c_*, *b_c_*, *h_c_* are respectively the apical length, basal length and height of the central cell without perturbing 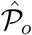 and *P_total_*. Moreover, the final tissue deformation was calculated as the relative extent of indentation of the apical and basal surface planes according to 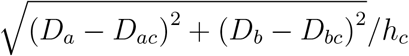 with *D_a/b_* being the fold depths of the apical/basal surface and *D_ac/bc_* being the apical/basal depth in the absence of perturbation.

## Appendix A Linking cell surface mechanics to cell and tissue shape via a 3D vertex model

Here, we sought to understand how epithelial tissue shape is determined by the mechanical properties of its constituent cells using the 3D vertex model described in Mathematical Modeling section. For a given set of cell mechanics, cell and tissue shape are derived from numerically computed vertex positions according to Eq. (6). The mechanically stable tissue shape is then obtained by minimizing the potential function *U*, as described in Eq. (7). Using this 3D vertex model, we studied the effect of locally induced modulation of cell mechanics on force-balanced tissue shape (Fig. S3A). For the shape stability analysis, see Appendix B.

Apical constriction of cells at the center of the tissue, highlighted in yellow, produces narrow apices, elongated lateral surface and widened basal membrane, and the tissue forms a shallow indentation (Fig. S3A, upper-left on the left panel). In contrast, basal expansion and lateral shrinkage induce laterally shortened and basally widened cells with minimal changes on the apical side, leading to deep, out-of-plane deformation (Fig. S3A, upper-left on the right panel). These results indicate that quantitative features of cell shape change and tissue deformation depend on the surfaces at which the mechanical modulation is applied, confirming our previous findings using a 2D model [41].

We next analyzed cell deformation in three conventionally categorized cell types, the columnar, cuboidal, and squamous cells (Fig. S3B), to better understand how cells respond to applied mechanical modulation. We found that in the columnar (◯) and cuboidal cells (△), changes in the apical and basal lengths, *a* and *b*, show negative correlation when apical mechanics 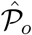 (Fig. S3B, left column) is altered. In contrast, changes in *a* and *b* are positively correlated in the squamous cells (□). This result indicates that different cell types show distinct deformation pattern in response to a given mechanical modulation. Furthermore, for a given cell type, cell deformation pattern depends on which cell surface is mechanically modulated. For example, for the columnar cells (◯), *a*–*b* and *a*–*h* are negatively correlated when apical mechanics is altered (Fig. S3B, left column), whereas *a*–*h* correlation becomes positive when basal cell mechanics *λ_b_* (Fig. S3B, middle column) is modulated, or *a*–*b* correlation becomes positive due to modulation of lateral mechanics *λ*_*ℓ*_ (Fig. S3B, right column).

Our 3D vertex model shows that different cell types or modes of mechanical modulation give rise to qualitatively distinct patterns of cell shape alteration, confirming our previous 2D analysis [41]. We suggested previously that experimentally derived correlation pattern on cell shape changes can be used to infer the mechanical principles that induce the shape changes [41]. Our current results suggest that this methodology can be extended to a 3D analysis.

## Appendix B Force balance analysis of 3D epithelial cells

According to Eq. (7), the potential energy function of an epithelial cell with the shape of a regular hexagonal prism reads,

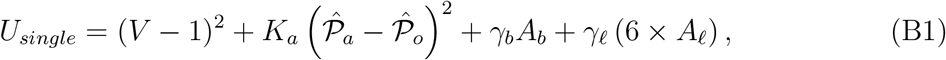

where 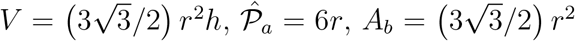, and *A_ℓ_* = *h* × *r* with *h* and *r* = *a* = *b* being the height and radius of the hexagonal prism, respectively (see Fig. S3B, lower right legend). From Eq. (B1), the force balance condition is given by the following relations:

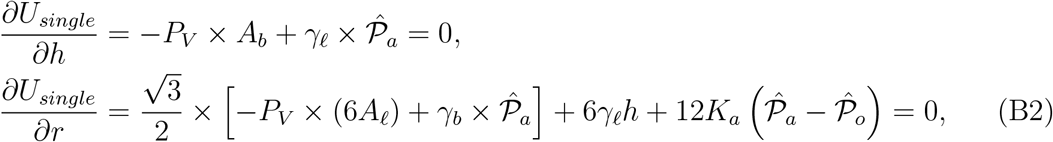

where *P_V_* = 2(1 − *V*) is the pressure resulting from cell volume elasticity. For a compressed cell (i.e., *P_V_* > 0), intracellular pressure pushes outward against shrinkage of compressive cell surfaces, and vice versa. To prevent cells from being distorted into frustum shape, we further considered the following condition:

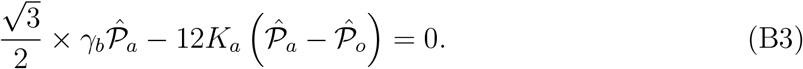

Therefore, from Eqs. (B2) and (B3), the condition for maintaining the shape of a hexagonal prism is

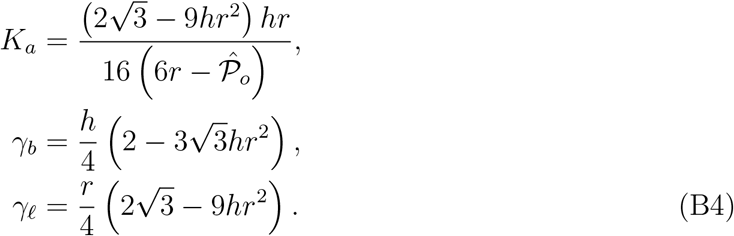

The stability of the cell shape can be determined from the Hessian matrix *M* of *U_single_* with the following components,

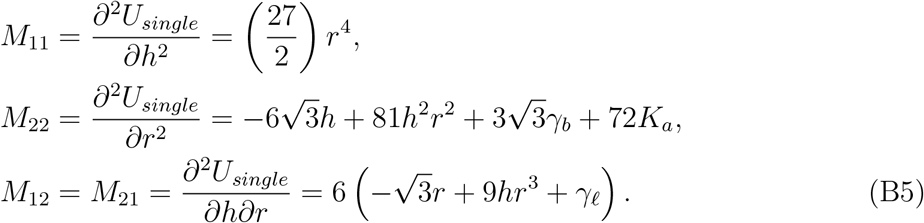

Cell shapes are mechanically stable when the trace and the determinant of *M* are positive, which gives the stable condition, as follows:

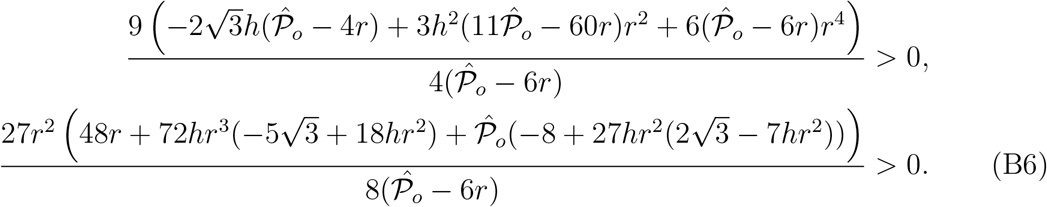

Note that *K_a_*, *γ_b_*, and *γ*_*ℓ*_ are replaced by *h*, *r*, and 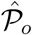 according to Eq. (B4).

## DATA AVAILABILITY

The data that support the findings of this study are available from the corresponding author upon reasonable request.

## AUTHOR CONTRIBUTIONS

F.L.W. and T.S. conceived the project; F.L.W. and T.S. performed the mathematical modeling; C.W.K. and Y.C.W. performed the imaging experiments and quantitation; All authors analyzed the theoretical and experimental data and participated in the writing of the manuscript.

## ACKNOWLEDGMENTS

We thank Chiao-Yu Tseng and members of the Shibata and Wang laboratories for valuable discussion and feedback. This work was supported by the core funding at RIKEN Center for Biosystems Dynamics Research; JSPS KAKENHI Grants JP19H00996 (T.S.) and JP18H02441 (Y.C.W.); JST CREST Grant JPMJCR1852 (T.S.); RIKEN Special Post-doctoral Researcher Program (F.L.W.).

---

**FIG. S1.**
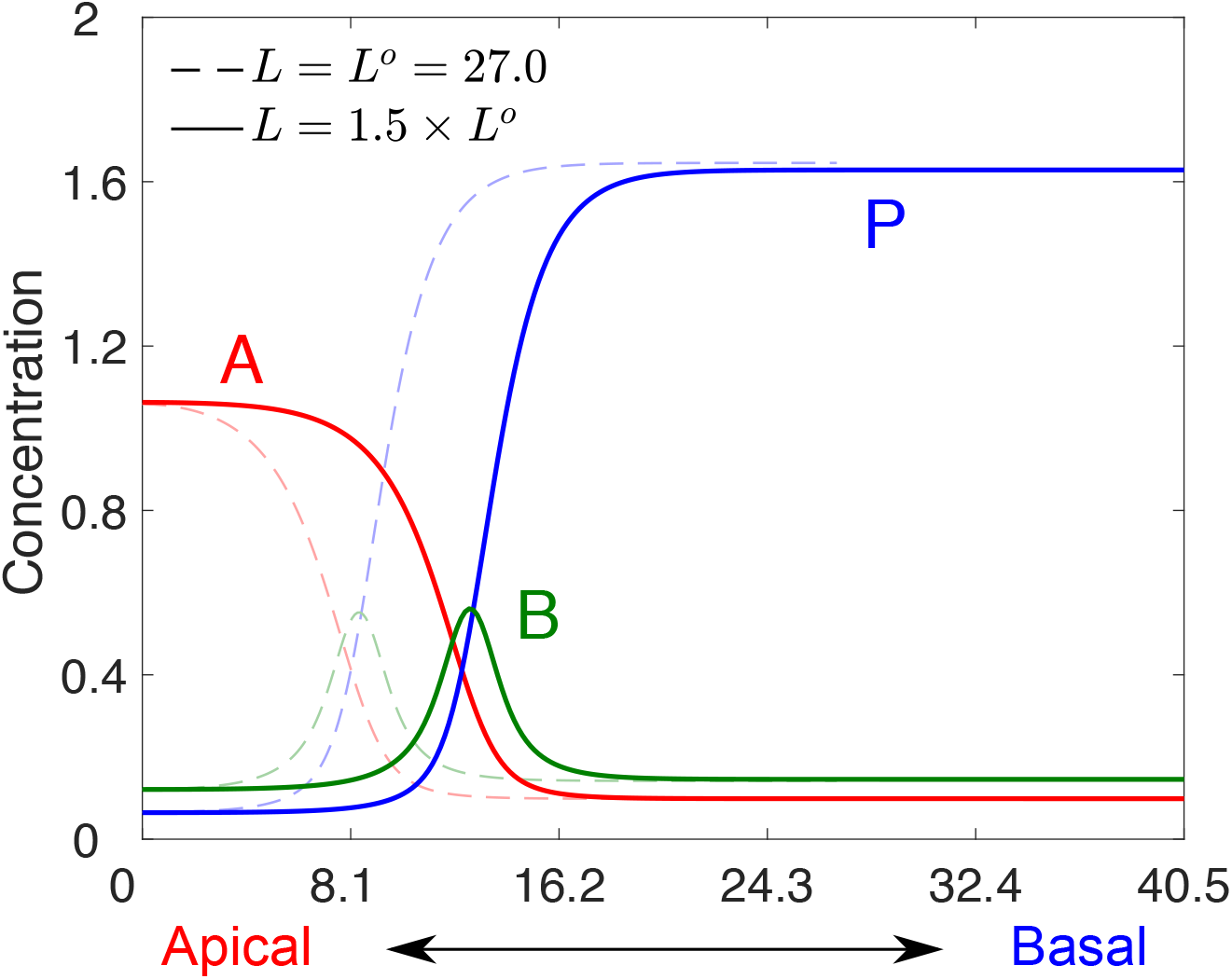
Basal shifts of the Bazooka peak position at the steady state (solid line) following an increase in *L* as compared to the control (dashed line). Note that the polarity distribution is shown on the absolute scale in contrast to Fig. 3B where the membrane location is scaled by *L*.

**FIG. S2.**
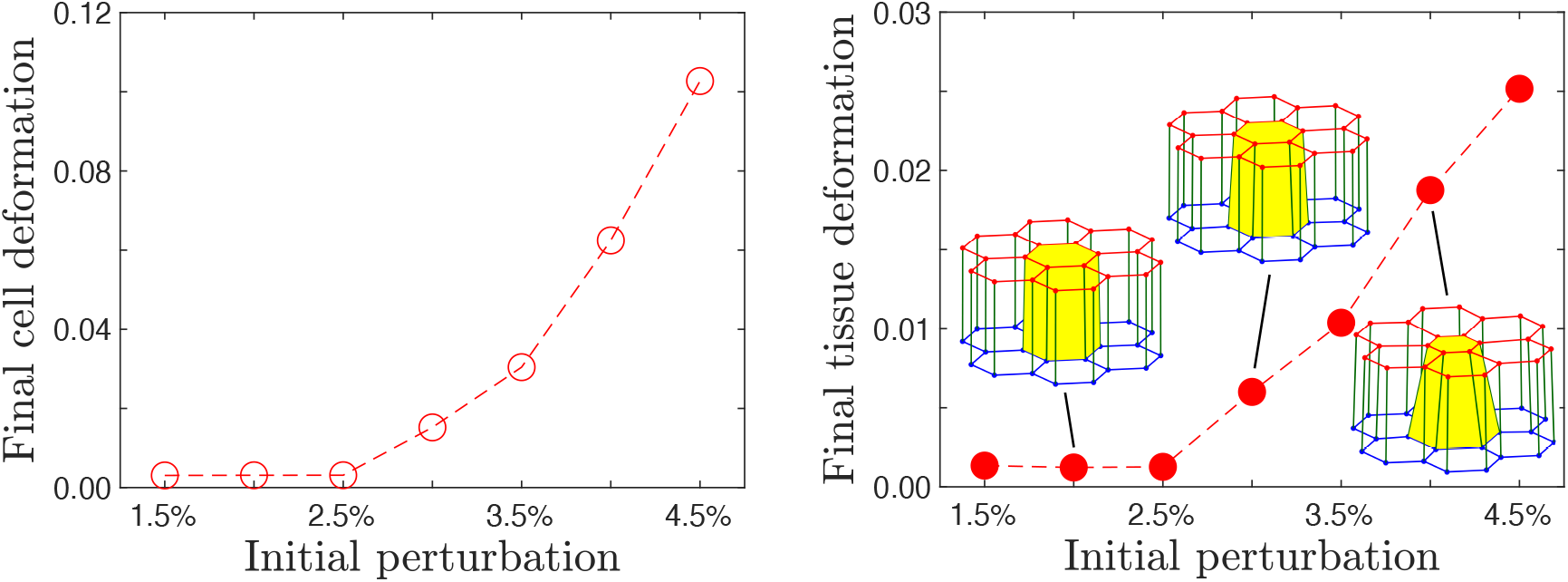
Quantitative analysis of biochemically triggered cell (left column) and tissue (right column) deformation under control of the mechano-polarity feedback. Insets in the right column show 3D visualization of the final steady-state tissue morphology following the initial biochemical perturbations. See Materials and methods for details of the analysis.

**FIG. S3.**
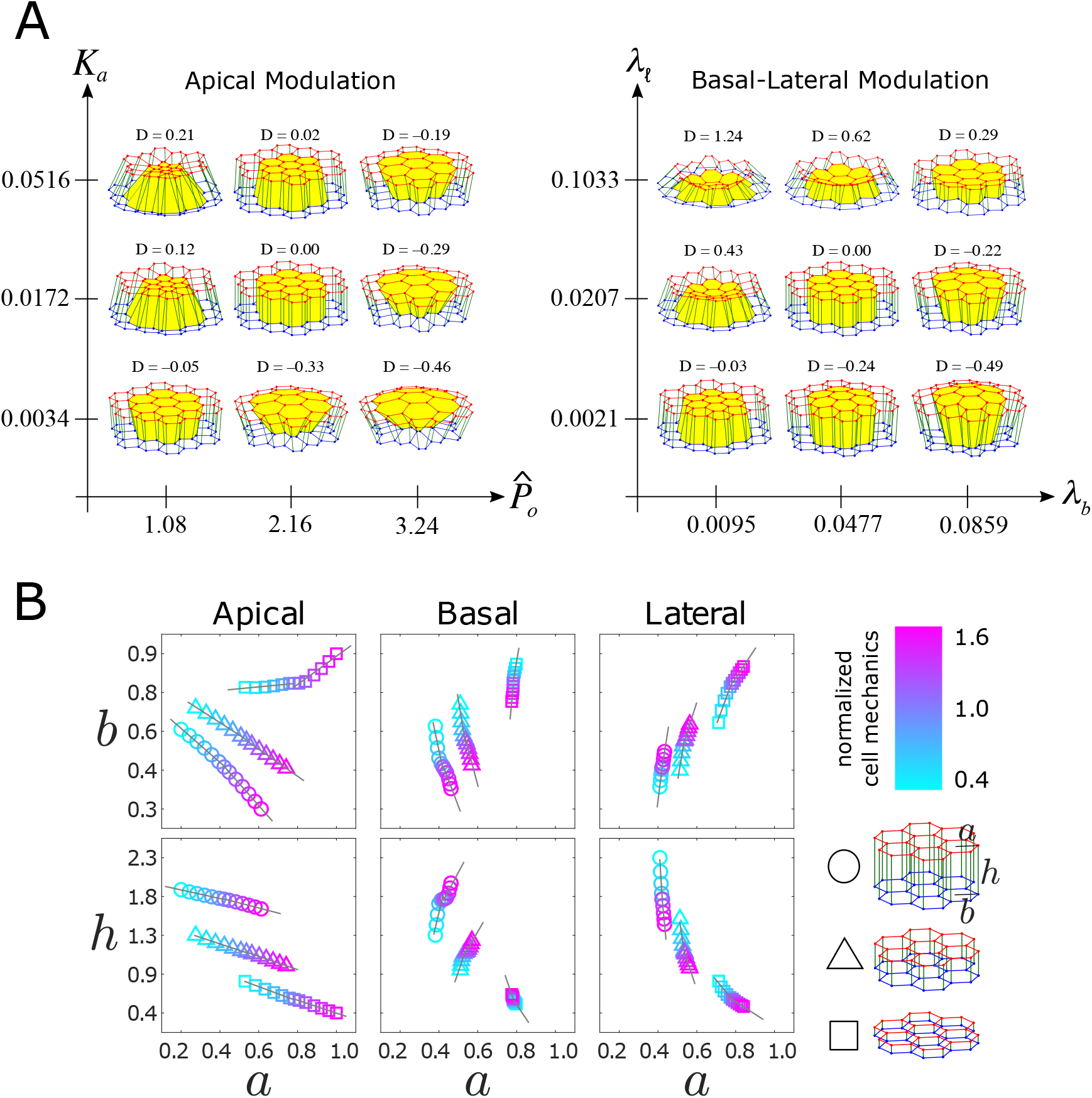
Epithelial deformation induced by modulation of mechanical parameters in a 3D vertex model. (A) A folded tissue structure forms when the apical mechanics, *K_a_* and 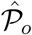 (left), or the basal– lateral mechanics, *λ_b_* and *λ*_*ℓ*_ (right), of a local cell population (yellow) is modulated. The fold depth *D* is calculated as the difference between the highest and the lowest vertices on the apical surface, where *D* > 0 (*D* < 0) indicates inward (outward) folding. (B) Cell deformation patterns in response to modulation of the apical (left column), basal (middle column), or lateral cell mechanics (right column). For columnar (◯; *a/h* = 0.25), cuboidal (△ *a/h* = 0.50), and squamous (□ *a/h* = 1.50) shapes, basal length *b* (top) and cell height *h* (bottom) are plotted against apical length *a* at the steady state for different values of mechanical parameters. The color bars indicate the relative deviation of parameter values from those of the flat epithelium, where 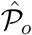, *λ_b_* and *λ*_*ℓ*_ are either increased or decreased for all the cells in the epithelium. The solid lines are linear regression lines that serve as a visual guide.

